# Model Agnostic Conditioning of Boltzmann Generators for Peptide Cyclization

**DOI:** 10.1101/2025.08.15.670426

**Authors:** Ben du Pont, Alex Abrudan, Rob Scrutton, Vladimir Radenkovic, Haowen Zhao, Tuomas Knowles

## Abstract

Macrocyclic peptides offer strong therapeutic potential due to their enhanced binding affinity and protease resistance, but their design remains a challenge due to limited structural data and tools that address only a narrow set of cyclization chemistries. Moreover, existing models are built to only consider ground state or mean conformations, rather than conformational ensembles that more accurately describes peptides. We introduce CycLOPS (a Cyclic Loss for the Optimization of Peptide Structures), a model-agnostic framework that conditions Boltzmann generators to sample valid cyclic conformations—*without retraining*. To overcome the scarcity of cyclic peptide data, we reformulate the design problem in terms of conditional sampling over linear peptide structures via chemically informed loss functions. CycLOPS encompasses 18 possible inter-amino acid crosslinks enabled by 6 diverse chemical reactions, and is readily extensible to many more. It leverages tetrahedral geometry constraints, using six interatomic distances to define a kernel density-estimated joint distribution from MD simulations. We demonstrate CycLOPS’s versatility via two distinct generative models: a modified Sequential Boltzmann Generator (SBG) (Tan et al., 2025) and the Equivariant Normalizing flow (ECNF) of Klein & Noé (2024). In both settings, CycLOPS successfully biases the Boltzmann distribution toward chemically plausible macrocycles.

## 1 Introduction

Macrocyclic peptides represent chains of amino acids modified through intramolecular crosslinking to form a topological loop and represent a promising class of therapeutics. Cyclization confers multiple advantages: resistance to denaturation under extreme conditions, reduced entropic cost of folding leading to increased binding affinity (Craik, 2006; Zorzi et al., 2017), protection from endogenous proteases (Hayes et al., 2021), and enhanced membrane permeability (Mizuno-Kaneko et al., 2023; Ji et al., 2024). These properties have established cyclic peptides across therapeutic domains, from cancer treatment (Ramadhani et al., 2022; Zhou et al., 2019; Mizuno-Kaneko et al., 2023) to antibiotics (Lai et al., 2022). As of May 2023, 46% of the 114 peptides approved in major pharmaceutical markets are macrocyclic (D’Aloisio et al., 2021; Costa et al., 2023).

Recently, there has been strong interest in the use of generative models for cyclic peptide design (Li et al., 2025; Rettie et al., 2025; Zhou et al., 2025; Zhu et al., 2025). Despite this, current computational design approaches exploit only a fraction of possible cyclization chemistries. Most methods are limited to head-to-tail or disulfide bridge cyclizations, overlooking the diverse chemical strategies known (Bechtler & Lamers, 2021). Even recent advances like CycDockAssem (Zhang et al., 2025) rely on individual bond constraints without considering the conformational flexibility of cyclic linkages. The work of Jiang et al. (2025) represents a significant advance in this area, decomposing all possible cyclizations into distance and type constraints on amino acids which can be learned from linear data alone. Yet their approach is limited in that it requires retraining for each additional cyclization added, making it unable to integrate with already well optimized models. Furthermore, they only condition on 1D pairwise Carbon-α distances between amino acids. This does not encode the required all-atom geometric constraints to enforce the formation of chemically valid cyclisations, let alone managing the modifications of chemical groups involved in the cyclisation reaction. More fundamentally, these techniques generate single average or ground state conformations rather than the ensemble of states p adopt in solution (Huang & Nau, 2003)—a critical limitation since such dynamics govern binding (Buch et al., 2011), and given the increasingly studied functionality of disorder (Holehouse & Kragelund, 2024).

Developments in neural generative modeling offer new possibilities. Flow matching and diffusion-based approaches (Lipman et al., 2022) have powered successful protein design tools like RFDiffusion (Watson et al., 2023) and structure prediction like AlphaFold (Jumper et al., 2021). Driven by a similar underlying technology, Boltzmann generators (BG) are a class of model which instead learn to sample from conformational distributions by mapping simple noise to approximate peptide structures (Noé et al., 2019). Such networks broadly fall into two categories: continuous normalizing flows (CNFs) and discrete normalizing flows (DNFs). The former are a special kind of neural ordinary differential equation (ODE)—that is, a network whose output is a velocity field

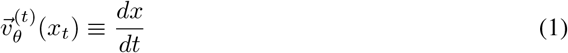

which represents the derivative of the noise w.r.t. which evolves samples from a prior along virtual time *t* (Chen et al., 2018). As these are typically applied to an easy to sample from prior, they can be thought of as inducing a continuous series of probability distributions, known as a flow, which interpolate between the prior and posterior.

Meanwhile, DNFs learn a fixed number N ∈ ℕ of subsequent invertible transformations that map a sample from a prior from an approximate posterior distribution; more formally, such a model may be defined as

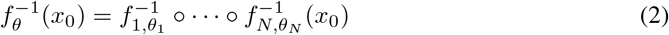

where x_0_ ∼ q(*x*), our prior, and θ_*n*_ represents the subset of our total parameters that determine that layer (Ho et al., 2019). The layers appear in reverse order and are written in terms of an inverse because of the way we must train such flows; given we know little about the data distribution that we hope to model, *p, a priori*, we must construct our loss in terms of the density of our prior. As such, θ is often found by attempting to minimize the negative log likelihood of our transformed observed data under our prior, which may be written as

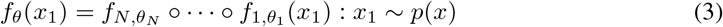

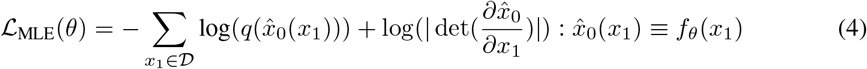

where *D* represents our observed samples from *p*, and 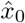 is approximately drawn from our prior (Ho et al., 2019; Zhai et al., 2024b). As Zhai et al. (2024a) note, this has an intuitive explanation: the first term in the sum attempts to bring data points close to the modes of our prior, whist the second term drives apart proximal datapoints, preventing model collapse. To clarify, there is no difference between the modalities upon which CNFs and DNFs operate. Rather, the latter conceptualizes its generation in terms of a finite number of steps, whilst the former defines an ODE which must be solved.

Both continuous (Klein & Noé, 2024) and discrete normalizing flows (Tan et al., 2025) have shown promise for efficient conformational Boltzmann sampling. However, conditional Boltzmanngenerator-based peptide design remains largely unexplored. The scarcity of cyclic peptide conformational data and its complicated chemistry, often involving non-canonical amino acids, have hampered the development of cyclization-conditioned peptide conformational generators.

**Our contributions:** This work introduces a Cyclic Loss for the Optimization of Peptide Structures (CycLOPS):

- CycLOPS is the first ready-to-use package to successfully condition Boltzmann generators for peptide design. It generates all-atom cyclic peptide conformations by conditioning the Boltzmann distribution.
- Our framework overcomes the lack of non-canonically modified peptide data by leveraging available MD simulations on canonical linear peptides together with chemically informed geometric constraints on 4 canonical atoms shared with the crosslink. We incorporate 18 cyclization strategies implemented via KDE-fitted tetrahedral conditioning, which is based on toy MD linkage simulations. This represents a significant innovation from previous work, since it simultaneously achieves strategy diversity and all-atom level conditioning.
- CycLOPS can condition any Boltzmann generator architecture, *without retraining* – we exemplify this by conditioning DNF and CNF models with simulated annealing and loss guidance, respectively.

## 2 Methods

### 2.1 Problem Formulation

A significant bottleneck in applying machine learning to science is the availability of large, highfidelity, and well-balanced datasets. This is also true of cyclic peptide design. Therefore, an essential question is how one can identify cyclizations of a linear chain that do not perturb its ensemble properties enough to affect its binding to a particular protein target. Moreover, can this be done without additional cyclic-peptide MD simulations or per-cyclization retraining? This may be reframed as a problem of statistical conditioning. First, we consider what a peptide’s linear conformations reveal about (1) its binding and (2) its possible cyclizations. (1) is well-studied in the case of structured targets. It has long been known that a protein’s structure is linked to its function. Hence, conformational motifs that are preserved across most of the conformational ensemble of the bound state play a crucial role in the binding interactions to a protein target. In fact, motif-constrained design has become a standard approach for developing protein binders (Yim et al., 2024; Ingraham et al., 2023; Trippe et al., 2022; Wang et al., 2022). Yet (2) remains largely underexplored.

Consider the random variables *S*, a possible conformation of a linear amino acid chain, and *S*_*κ*_, a possible cyclic conformation in any possible linkage, both implicitly conditioned on a particular initial sequence of amino acids. Of course, the distribution of *S*_*κ*_ is not completely knowable given the probability density function of *S* alone, since the chain never truly exists in a cyclic conformation; at no point do any of the bonds involved exist. Therefore, we approximate this as a problem of statistical conditioning.

Intuitively, nearly cyclic conformations—structures which almost satisfy typical bonding constraints—appearing with significant probability density suggest that the linear chain may be amenable to a given cyclization. Let *C* be the event the chain is approximately cyclic, i.e. the constraints of a particular linkage are almost satisfied. The problem of cyclic peptide design then becomes sampling from valid linear conformations conditioned on both binding feasibility and cyclizability, as shown in the inner purple region of Fig. 2. We therefore seek to modulate the landscape of *S* to increase the probability of sampling approximately cyclic peptides, ideally whilst preserving the dynamics of a binding region or motif. To sufficiently limit the scope of this paper, we shall principally consider *S*|*C*, as conditioning on binding is somewhat well studied.

**Figure 1.**
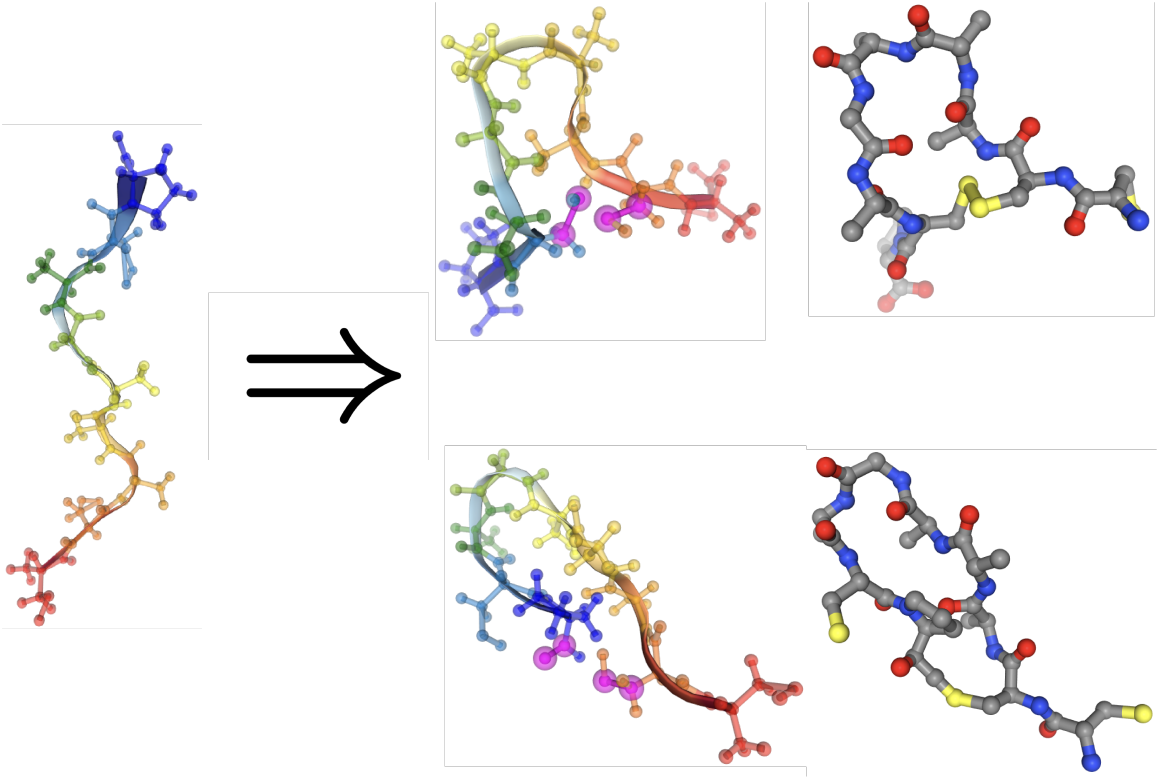
Cyclization of a linear peptide at two different stapling sites performed through CycLOPS conditioning of a Boltzmann Generator.

**Figure 2.**
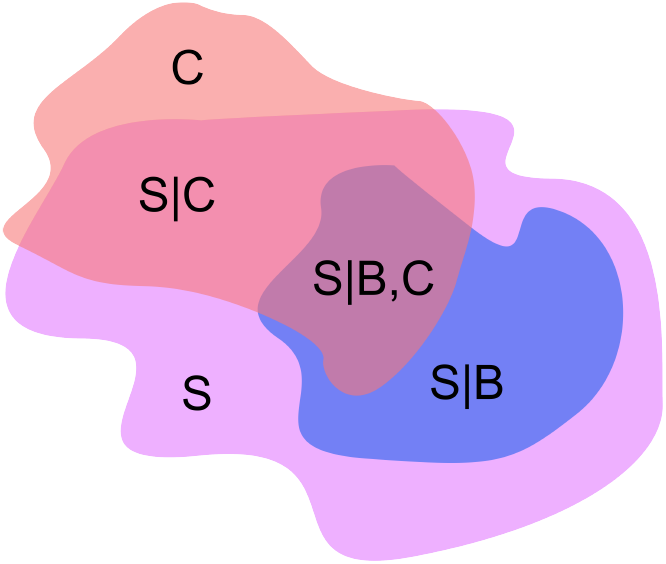
Schematic representation of peptide conformational spaces showing linear Boltzmann space (*S*), cyclic space (*C*), and bound (*B*) regions, with *S*|*B, C* representing the desired sampling target. Note that *S*|*B* is equivalent to *B*, because a state can only be bound if it is already a valid conformation of the chain. Similarly, it is trivial to arrive at cyclizations which completely perturb a linear chain and whose constraints are never approximately satisfied. Thus *C* is not a subset of *S*.

### 2.2 Tetrahedral Geometry for Cyclization Constraints

How might one restrict a chain to exist only in approximately cyclic conformations? A natural approach uses *constraint dissolving functions* (CDFs)—functions whose optima satisfy the desired condition. However, simple distance or 2D angle constraints fail to capture the conformational flexibility of complex linkages involving multiple atoms. But how can we score how well our chain’s positions respect a given linkage’s constraints? Ludwig Boltzmann’s studies of the micro-canonical ensemble show that

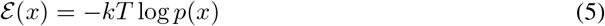

This gives us an excellent CDF whose minima represent modes of our probability distribution.^1^ Our insight is that any linkage can be characterized by four atoms it shares with the original peptide chain. These four atoms define a tetrahedron, and the six distances between them fully specify the spatial relationships required for valid cyclization, given a sufficiently small linkage.^2^ Let **x** = [d_01_, d_02_, d_03_, d_12_, d_13_, d_23_] be the ordered vector of these edge distances. This representation preserves *E*(3) invariance that is essential for maintaining the geometric properties of our Boltzmann generators. To estimate the probability of a chain’s positions under a given cyclization, we then isolate the linkage in question and simulate it via Langevin molecular dynamics (MD). By construction, these small molecules are toy models of the linkage that share at least four atoms with the original chain. We assume the linkage is sufficiently small to avoid forming non-physical clashes with the rest of the peptide. As such, if we can compute the probability density of the relative positions of these points under the toy cyclization model, we can approximate the probability density of the entire chain’s conformation *x* under that particular cyclization, and thus ℰ (*x*) by Eqn. 5.

Yet, MD simulations yield collections of points, not probability densities. One approach to circumvent this is to use generative models, which have become increasingly popular for density estimation (Ho et al., 2019; Dinh et al., 2016). However, these are slow, prohibitively so for our purposes. As such, we employ a much older technique, Kernel Density Estimation (KDE) (Rosenblatt, 1956; Parzen, 1962), to convert our data to approximate distributions:

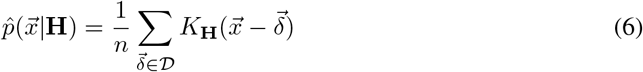

And

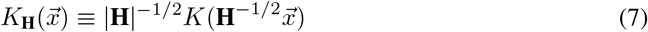

where K is our kernel, *D* contains our simulated tetrahedral distances for the linkage in question, **H** represents a positive definite bandwidth matrix and **M** denotes the determinant of matrix **M**. In this instance, a kernel is a positive function which integrates to one.

For ease of optimization, we additionally require K be unimodal, spherically symmetric, centered on the origin, and smooth (Chacón & Duong, 2018), with heavy tails to ensure finite values once the log is applied and non zero gradients far from the data. Thus the Cauchy distribution

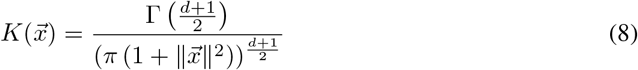

is ideal for the reasons it is normally difficult to work with.

### 2.3 Multi-Cyclization via Maximum Entropy

Real peptides often admit multiple cyclizations, as illustrated in Fig. 3. Given individual CDFs ℰ_*i*_(*x*) for each possibility, how do we construct a valid overall cyclic loss? We desire an expected cyclic “energy” given the chain is cyclic:

**Figure 3.**
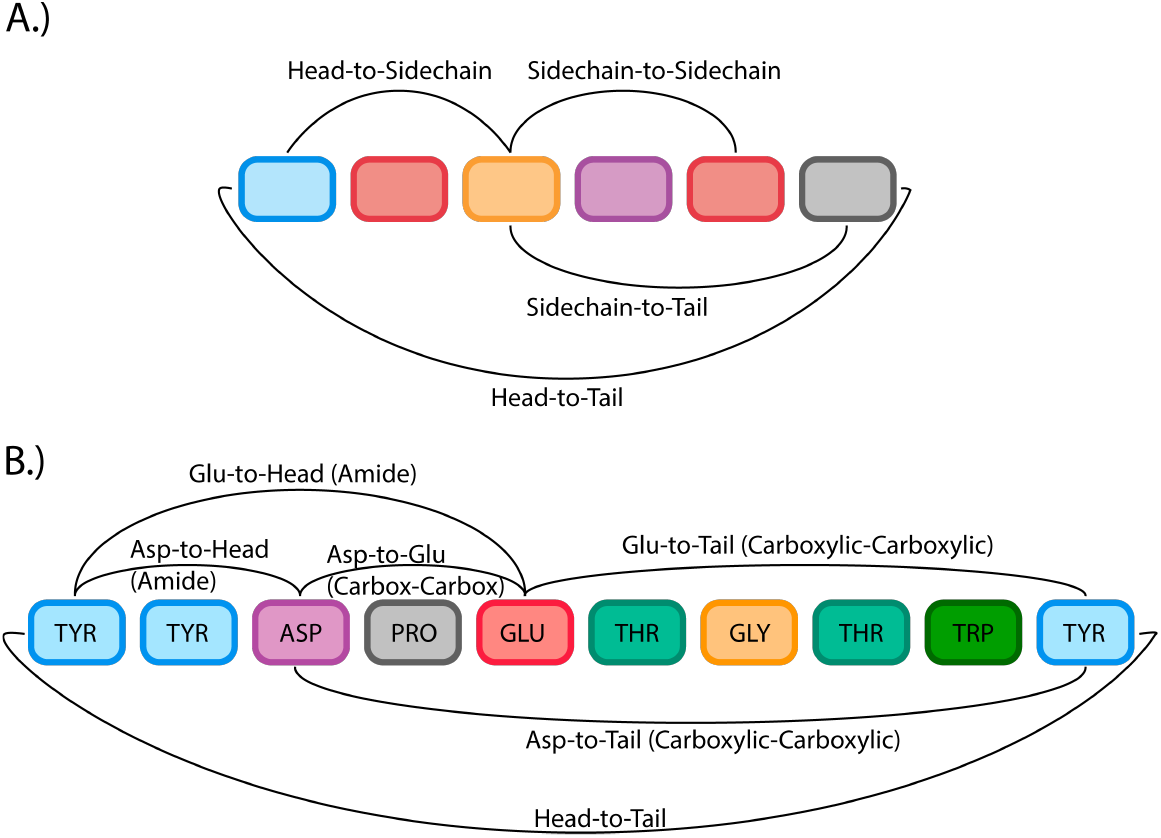
**A.)** A diagram illustrating some of the chemical permutations possible for peptide cyclizations. **B.)** Some of the possible cyclizations for the chignolin peptide (Suenaga et al., 2007), based on some of the chemistries from Bechtler & Lamers (2021).

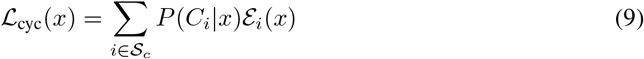

where *S*_*c*_ contains all cyclization strategies and *P*(*C*_*i*_|*x*) is the probability of cyclization i given conformation *x*. A naive way to accomplish this is simply to assume a uniform reweighting:

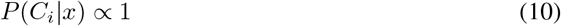

Yet this takes nothing into account about what we know about each cyclization’s relative energy, and may guide the system into unphysical superpositions of cyclic states. Instead, we weight by the relative probabilities:

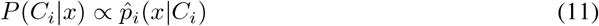

By Eqn. 5, this becomes:

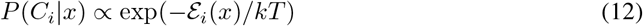

This Boltzmann reweighting is maximally entropic—it assumes the least about the underlying cyclization probabilities (the standard exponential result doesn’t apply here since we’re computing, not constraining, the expectation; see Appendix A.1 for a quick proof of the claim). By decoupling the reweighting temperature *T*_*d*_ from the individual energy temperatures, we define:

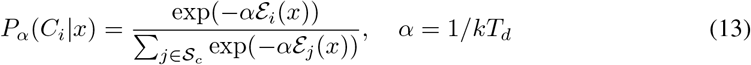

As α → ∞, this approaches a hard minimum, preventing optimal structures from being unphysical superpositions. Thus we define:

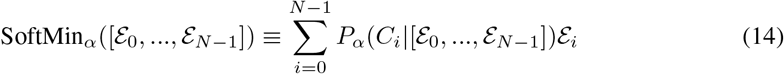

Interestingly, this is tantamount to renormalizing, given a set of energies, according to what their probabilities *would have been* if generated with temperature *T*_*d*_.^3^

For resonant cyclizations (composed of sets of chemically equivalent atoms), multiple losses may represent the same chemistry. Since these are geometrically similar, including them all in the Soft-Min may locally approximate an average, again yielding non-physical constraints. We wrap resonant losses in a hard minimum before applying SoftMin to avoid this. Another intuitive way of looking at this is that these two cyclizations are physically indistinguishable, and therefore the same. It would thus be unideal to “doubly” count them. This and the rest of the CycLOPS framework is illustrated in Fig. 4.

**Figure 4.**
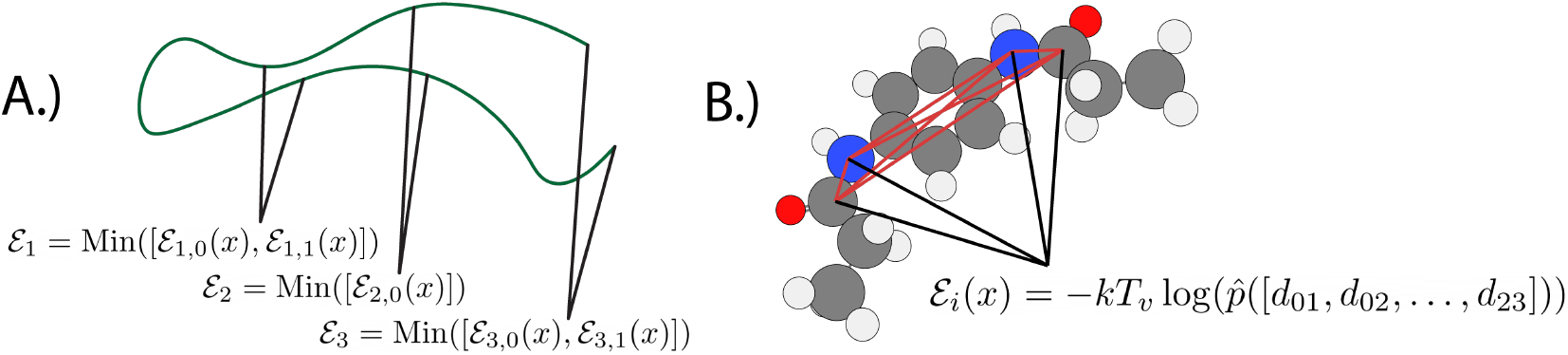
Schematic representation of the CycLOPS framework, showing **A.)** the automatic recognition of diverse possible cyclization sites on a peptide chain and **B.)** the computation of individual cyclization CDF values via a tetrahedral KDE fit on simulations of a toy model and evaluated on the four atoms in common with the original chain.

### 2.4 Model-Agnostic Conditioning

Given our cyclic loss ℒ_cyc_, how do we condition Boltzmann generators without retraining? A universal method is to optimize over model latent space, searching for conformations which minimize our CDF (Abdin & Kim, 2024; Noé et al., 2019). If the underlying network is differentiable, one could use a stochastic gradient descent-based optimizer, like ADAM (Kingma & Ba, 2014). A strategy without this requirement, however, is *simulated annealing*, which has already seen success in cyclic peptide design (Zhu et al., 2025). This is a probabilistic optimization algorithm inspired by annealing in metallurgy, where a material is heated and then slowly cooled to reach an energetic minimum (a highly ordered crystal structure). Here, our system’s “energy” is some function of our state we seek to minimize. As illustrated in Alg. 1, simulated annealing involves sequentially updating some state *x*_*t*_ by proposing a new state *x*^*′*^, computing the difference between their energies, and switching states as a function of some gradually cooling temperature and the change in energy. As the temperature cools, the algorithm gets more greedy, therefore modulating between exploratory and exploitative behavior as it runs its course. Thus it is not as prone to local minima as other optimization algorithms if well tuned (Press et al., 2002).

#### Algorithm 1

Generic Simulated Annealing

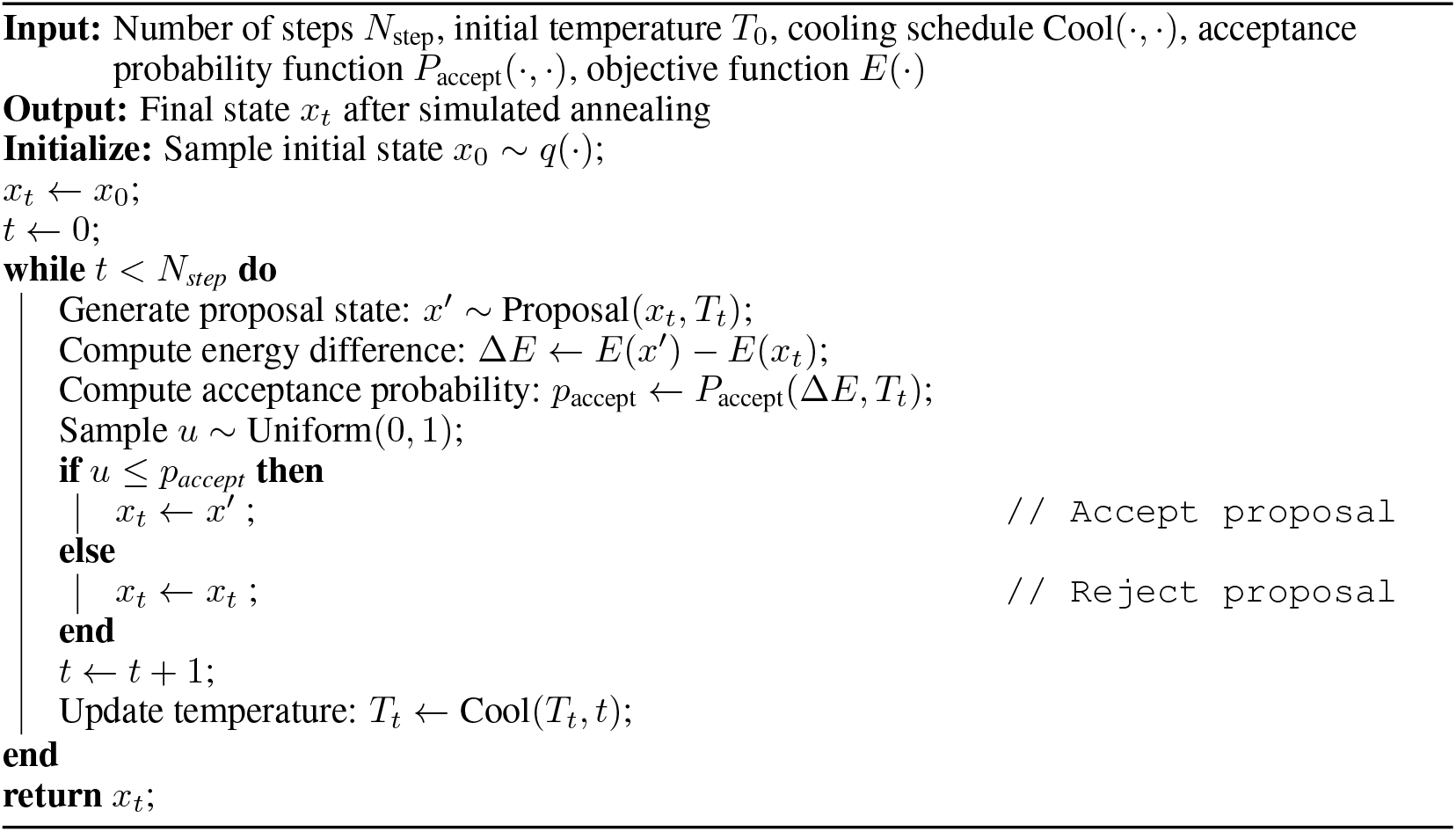

Simulated annealing is, by nature, suited to discrete optimization. We adapt this to our continuous usecase by assigning the system a velocity *r*_*t*_ at each timestep, which then determines the radius of a hypersphere from which a proposal state is sampled from. Inspired by the relationship between temperature and mean particle velocity, we allow *r*_*t*_ to scale with the square root of the ratio of the current to initial temperature. Since the probability of switching between states depends directly on Δ*E*/*T*_*t*_, care must be taken to pick an appropriate temperature; too large, and the system will “melt,” switching between states with no regard for their energy. Too small, and the system will collapse into whatever minimum is most proximal to its initial state. We therefore set an appropriate *T*_0_ based on a calculated mean absolute initial Δ*E*, which is then scaled by a user set hyperparameter κ that determines the starting switching probability. We then exponentially cool the simulation to zero by multiplying the previous temperature at each timestep by a constant λ ⪅ 1 to determine its new temperature; this is so that the system is cooled sufficiently slowly, which, given enough steps, helps the algorithm find a global minimum. This is elaborated upon more formally in Alg. 2 of Appendix A.2.

For continuous normalizing flows, direct latent optimization can be prohibitively slow. Instead, one can guide a CNF to sample from an approximate conditional posterior. In its simplest incarnation, this takes the form of Bayes rule (Chung et al., 2022; Jiang et al., 2025):

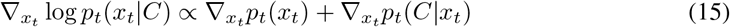

In many instances, the connection between *x*_*t*_ and *C* must be learned, which can be challenging; in these cases, it is preferable to construct a differentiable CDF for the condition in question Song et al. (2023). For denoising diffusion probabilistic models Ho et al. (2020)–since CDFs are naturally defined over the final structure space rather than the noisy intermediate states–we can employ the following approximation

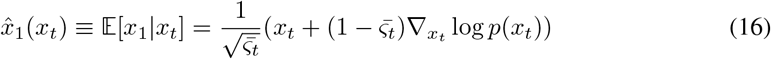

where *ς* : *t* ∈ [0, 1] → (0, 1] is some scheduler, 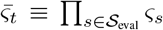 and *S*_eval_ is the set of discrete evaluation timesteps in (0, 1) (Ho et al., 2020; Komorowska et al., 2025). This has the intuitive explanation of linearly interpolating between the current state and the final state based on the present vector field of the neural ODE to arrive at an expected output. Thus, we get:

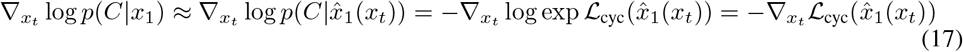

where ℒ _cyc_ enforces *C* at *t* = 1. Since diffusion is effectively optimal transport flow matching (Gao et al., 2024),^4^ we construct the heuristics:

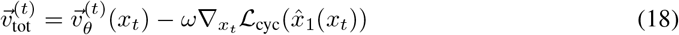

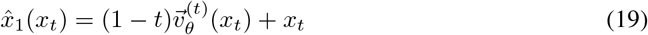

where ω > 0 represents our conditioning strength; greater values will help ensure the condition is satisfied, but will contribute to lower sample quality. Additionally, by the chain rule:

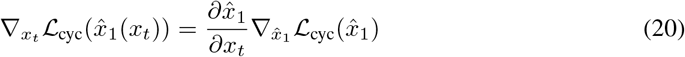

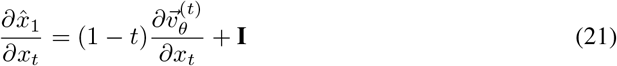

Komorowska et al. (2025) *note that this comprises of an easy to compute gradient* 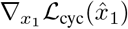 and an expensive Jacobian matrix 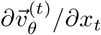, which can be empirically ignored (set to **0**). Thus, CycLOPS provides a unified framework for conditioning any all-atom Boltzmann generator toward diverse cyclic peptide conformations without retraining. This can be done in all cases via simulated-annealing-based latent space optimization, or flow guidance if working with a CNF.

### 2.5 Toy Simulations of Linkages

As illustrated in Fig. 4, CycLOPS relies on a knowledge of the approximate joint distributions of the edge lengths of a tetrahedron which encodes the geometry of a given linkage. We must therefore simulate the toy models of each of these linkages. In all cases, this was performed with the OpenMM python package (Eastman et al., 2013) with initial configurations generated via PACKMOL Martínez et al. (2009). Forces were generated via the Amberff14SB forcefield (Maier et al., 2015) and the TIP3 water model (Jorgensen et al., 1983). Molecules were parameterized via SMIRNOFF (Mobley et al., 2018) using OpenFF (Qiu et al., 2021; Boothroyd et al., 2023), with volume calculations handled by CCTK (Wagen & Kwan, 2020). Simulation code is adapted from Wagen (2024). Each small molecule was simulated for 5 ns on one A100-80GB VRAM GPU over 10 seeds with a step size of 1 fs at 300K and with a friction coefficient of 1 ps^*−*1^. The first nanosecond of each simulation was discarded for equilibriation, leaving a total effective simulation time of 40 ns. The molecule used to model each linkage’s constraints is shown in Fig. 5, along with the atoms used to define the tetrahedra on which the KDEs are fit. Given the smallness of each system, we observe that this was quite computationally inexpensive.

**Figure 5.**
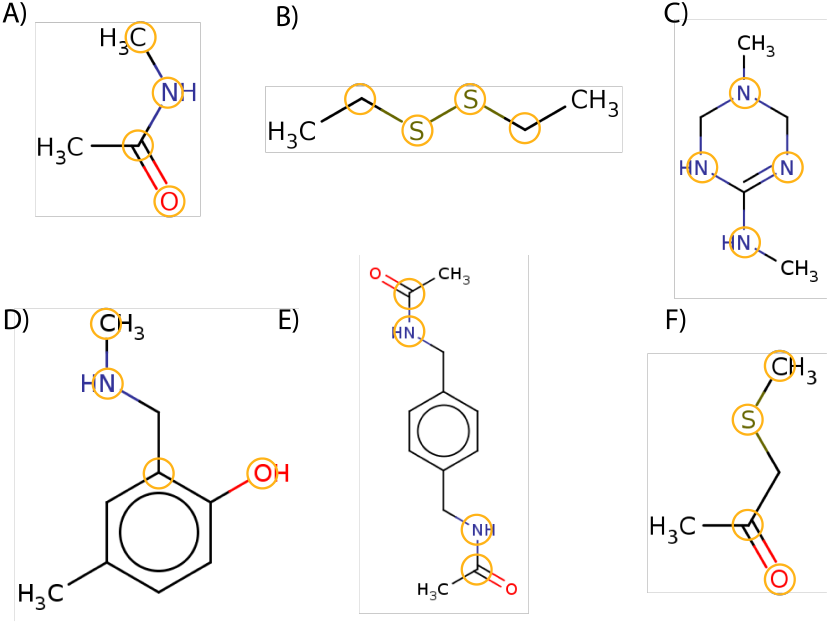
Schematic representations of the toy models used to fit KDEs for each cyclization chemistry. Shown are: **A)** amide bond cyclization model, **B)** disulfide bond cyclization model, **C)** lysine– arginine cyclization model, **D)** lysine–tyrosine cyclization model, **E)** carboxyl–carboxyl cyclization model, and **F)** cystine–carboxyl cyclization model. The atoms used to fit the tetrahedral KDE are highlighted in yellow. Note that these need not correspond to atoms of the same species after the chemistry is applied, highlighting the versatility of CycLOPS. For instance, the nitrogens of **E)** initially corresponded to carboxyl oxygens.

**Figure 6.**
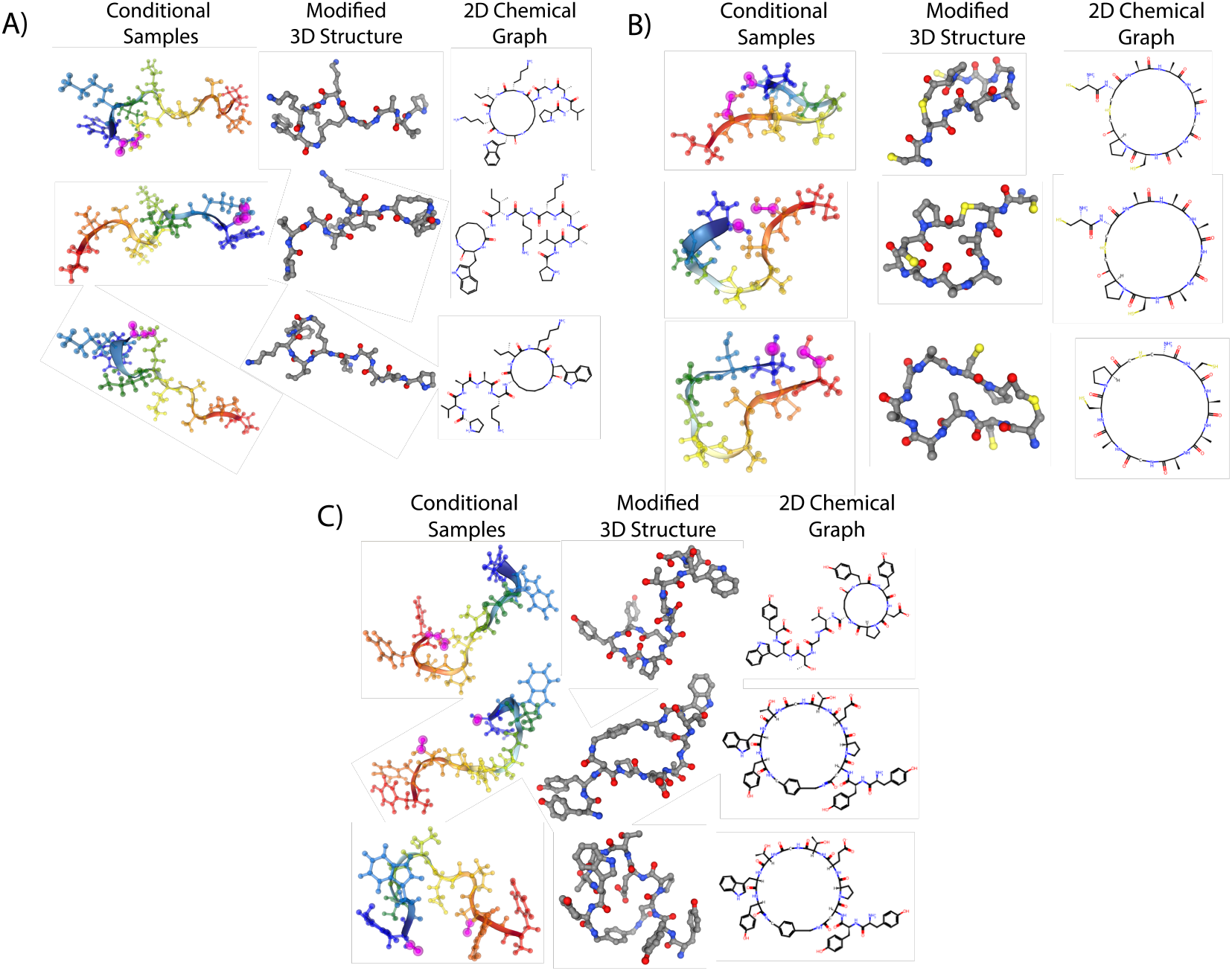
For three peptides **(A)** PVAAKKIKW, **(B)** CCAAAGACP, and **(C)** Chignolin: (*left*) select conditional samples generated via simulated annealing of the CycLOPS loss using the respective DNFs shown in Table 1, with pink highlighted atoms showing the cyclization site, (*center*) 3D models of said predicted cyclizations with cyclization atoms added (hydrogens omitted, only newly added atoms relaxed), and (*right*) 2D skeletal representations showing chosen cyclization linkages.

The joint distributions of the six distances defined by each of the four specified atoms were then fit with a KDE with a bandwidth of 0.64Å ∗ **I**. Ideally, contemporary multivariate kernel smoothing techniques, like the multivariate extension of the method of Sheather & Jones (1991) proposed by Chacón & Duong (2010), should be used; due to the non-triviality of developing a Python implementation, we leave this to future work. For PyTorch compatible kernel density estimation, such that loss calculations are automatically differentiable and benefit from GPU acceleration, we use the implementation provided by Kladny (2025). However, CycLOPS should not only compute ℒ_cyc_(*x*), but also seamlessly handle the identification of possible cyclizations. This functionality is implemented largely through the MDTraj Python package (McGibbon et al., 2015).

### 2.6 Model Training

We approximately follow the training procedures enumerated in Tan et al. (2025) to train our models, except without energy *W*_1_ distance based early stopping and without restricting our training set to the first 100,000 frames present. This is because we do not perform importance based annealing post generation, so our underlying model must be more robust. As such, we reduce the maximum number of epochs depending on the number of samples. We exclusively employ Tan et al. (2025)’s Chignolin TarFlow (Zhai et al., 2024a) architecture to study DNFs, and ECNF++ to study CNFs.

As large molecular dynamics trajectories are very expensive to simulate, we use those provided in the literature wherever possible. This notably includes *ab initio* Chignolin at DFT level (Wang et al., 2023), classical MD trajectories across diverse sequences (Zhu, 2021), and alanine peptides of various lengths (Schopmans & Friederich, 2025). Training and architecture specifics are provided in Table 1.

**Table 1:** Model Architectures and Training Details.

| Type | Sequence | T(K) | Trainset Size(Frames) | Data Src. | Params | Epochs |
| --- | --- | --- | --- | --- | --- | --- |
| SBG | GYDPETGTWG<br>(Chignolin) | 340 | 4.6M | Wang et al. (2023) | 114M | 80 |
| SBG | PVAAKKIKW | 300 | 480K | Zhu (2021) | 114M | 300 |
| SBG | CCAAAGACP | 300 | 480K | Zhu (2021) | 114M | 300 |
| ECNF++ | AAAA | 300 | 10M | Schopmans & Friederich (2025) | 2.3M | 10 |
| ECNF++ | AAAAAA | 300 | 10M | Schopmans & Friederich (2025) | 2.3M | 10 |
| ECNF++ | WLALL | 300 | 100K | Scherer et al. (2015) | 2.3M | 1000 |

**Table 2:** Training Details and Computational Resources.

| Type | Sequence | Batch Size (cumulative) | Number of GPUs | Total GPU Hours |
| --- | --- | --- | --- | --- |
| SBG | GYDPETGTWG<br>(Chignolin) | 2048 | 8 | 351.2 |
| SBG | PVAAKKIKW | 2048 | 8 | 304.6 |
| SBG | CCAAAGACP | 1024 | 4 | 241.2 |
| ECNF++ | AAAA | 2048 | 4 | 35.88 |
| ECNF++ | AAAAAA | 1024 | 4 | 81.2 |
| ECNF++ | WLALL | 1024 | 8 | 165.6 |

## 3 Results

### 3.1 Discrete Normalizing Flow Simulated Latent Annealing

As shown in Fig. 3.1, simulated annealing of the CycLOPS loss can produce valid cyclic structures given a Boltzmann generator trained only on linear peptide data. This appears to be true across most peptide sequence space, as evidenced by the three distinct sequences employed to generate subfigures A), B), and C). Moreover, the conditional samples shown in the first column of each subfigure correspond to valid 3D structures, as shown in the second column of each subfigure. These are generated by adding the atoms necessary for the chosen cyclization to the peptide with the correct bond topology, whilst deleting molecular fragments that are removed as reaction by-products. The atoms added, and only these atoms, are then relaxed as much as possible towards valid structures without unfairly advantaging our conditioning framework. It should be noted, however, that good initial guesses should be provided as the positions of these atoms because force field gradient descent, which is typically how relaxation is performed, is prone to getting trapped in local minima. Such minimizations are common in the context of MD, where pre-simulation minimization is typically designed to prevent large initial forces from crashing the protocol. Yet this is not sufficient for our purposes, so manual initial guesses must be made. In the rightmost column of each subfigure is a 2D representation of the chemistry used, further clarifying the location of the macrocycle. In the above relaxation protocol, the MDAnalysis Michaud-Agrawal et al. (2011); Gowers et al. (2016) Python package is used to parse peptide information from the conformation files. All relaxation, bonding and 2D graph generation is handled by the RDKit (Landrum et al., 2025) package. All 3D atomic visualizations are created via NGLView Nguyen et al. (2018). Samples shown in Fig. 3.1 based on visual inspection.

We examine not only individual samples produced by DNF CycLOPS conditioning, but also the effects this has on sample distributions. As shown in Fig. 7, for all peptides used, CycLOPS annealing guides samples toward statistically significant lower values of ℒ_cyc_. It should be noted that most of these samples do not correspond to valid cyclic peptides: in practice, we find that only about 10% of samples in each case do not have clashes. This sampling paradigm—generating many candidates and selecting the best—is ubiquitous across protein generative AI, where the power lies not in perfect individual samples but in the ability to rapidly explore chemical space and identify top-performing designs. Much of this, we observe, has to do with the aforementioned assumption of sufficiently small linkage chemistries being violated. For example, the inter-carboxyl benzene-ring-based chemistry shown in Fig. 5 E) is large enough to interact with atoms of the peptide chain, which is capped by the tetrahedral vertices. Unfortunately, we are unaware of an elegant solution to this problem; the addition of virtual atoms to a model’s chemical graph is likely to cause significant issues, since each network has only learned a singular topology. Furthermore, no universal Boltzmann generators have been released yet, and even such a model would be unlikely to be trained on non-canonical amino acids that are needed to form cyclic linkages. As such, we adhere to the previously described approach: generate many samples and eliminate the problematic ones. Clashes are also observed, however less frequently, in regions far from the linkage since conditioning drives latents somewhat out of distribution by definition.

**Figure 7.**
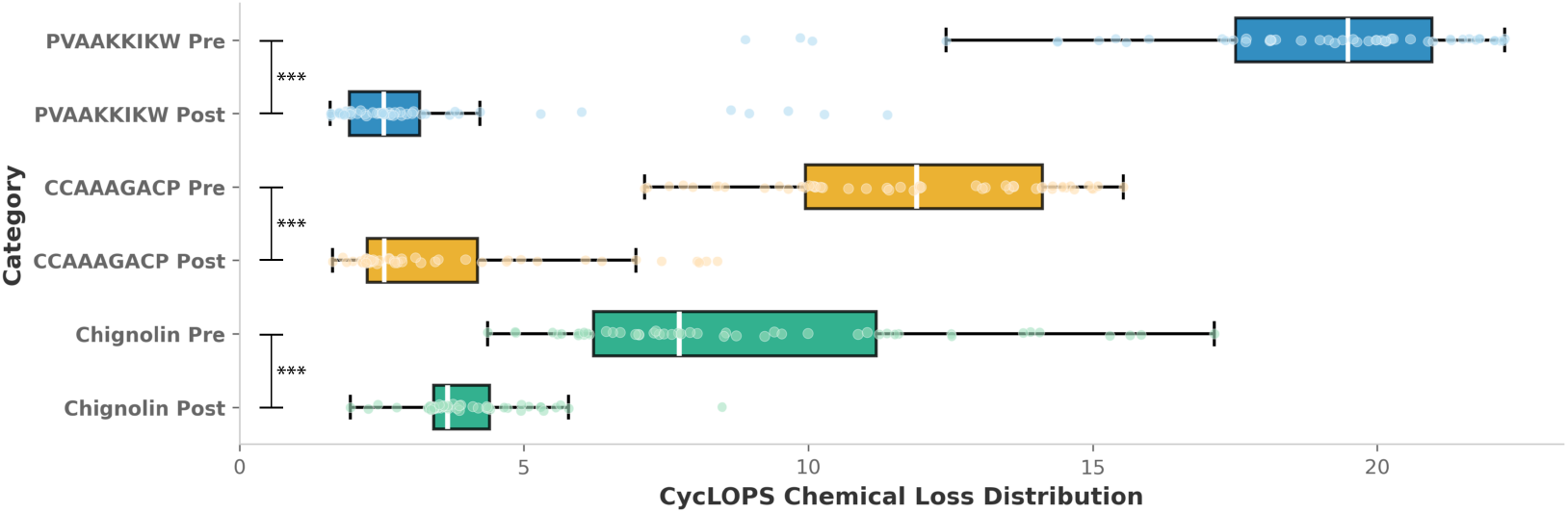
Box and whisker plot of the distribution of the CycLOPS cyclic losses of 50 random samples before and after simulated annealing for representative peptides. In all cases, simulated annealing significantly changes the distribution (*p* < 0.001 represented by ***, two-sample Kolmogorov– Smirnov test). All mean-to-mean differences between pre- and post-annealing distributions are significant up to 3 decimal places under a permutation test with 10,000 random permutations.

As seen in Fig. 8, DNF-simulated annealing appears to explore prior space since the distribution of finally chosen cyclizations differs from that of the smallest loss before annealing. In the case of PVAAKKIKW and CCAAAGACP, this involves changing the most frequent smallest loss. This is also apparent on a sample-wise level in Fig. 9, which shows the pre- and post-conditioning structures of the samples from Fig. 3.1. This suggests that the loss minimization observed in Fig. 7 results from genuine latent space exploration rather than simply refining the initially generated structure.

**Figure 8.**
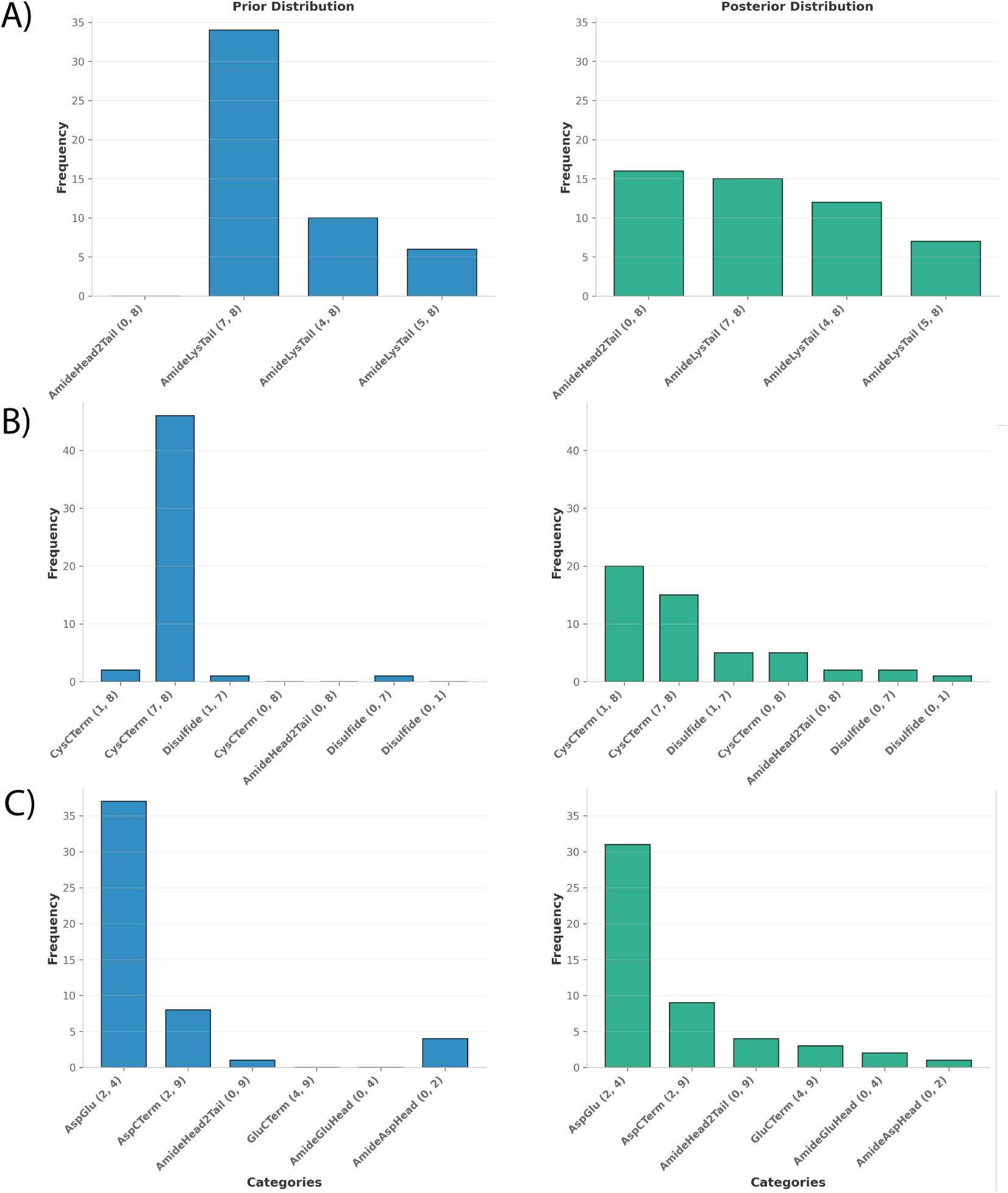
Prior and posterior distributions of minimum chemical losses reveal that cyclization preference shifts after simulated annealing. Histograms illustrate the lowest individual chemical loss within ℒ_cyc_ before (prior) and after (posterior) simulated annealing of 50 samples for **(A)** PVAAKKIKW, **(B)** CCAAAGACP, and **(C)** Chignolin. Each cyclization is labeled by its chemistry, with bonded amino acid positions in parentheses. Simulated annealing systematically shifts the loss distributions, with the most favorable cyclization changing for peptides A and B.

**Figure 9.**
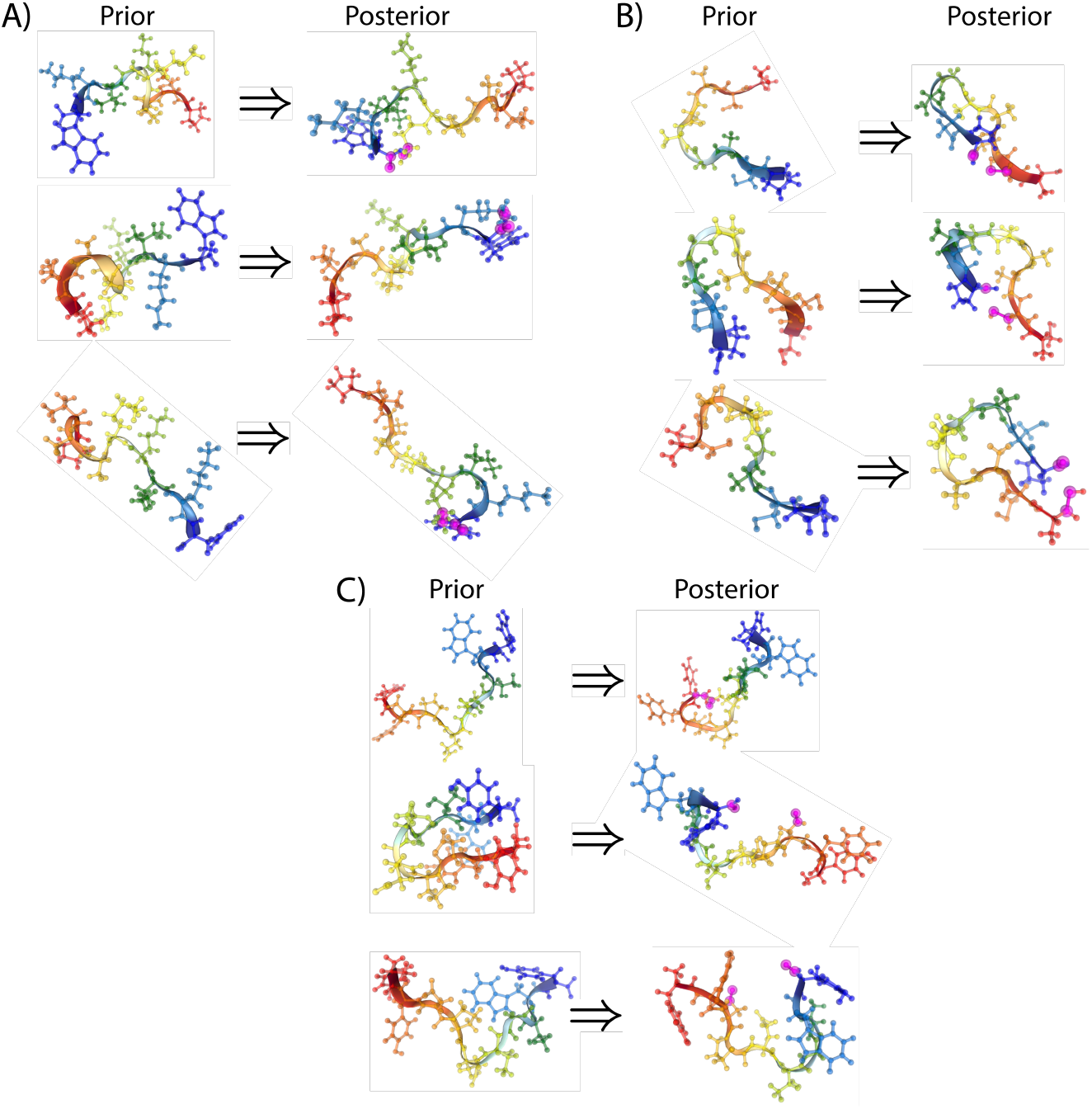
Apparent conformational changes in peptide samples from Fig. 3.1 before and after ℒ_cyc_ simulated latent annealing demonstrate latent space exploration. Sequences: **(A)** PVAAKKIKW, **(B)** CCAAAGACP, **(C)** Chignolin.

The CycLOPS framework trivially enables specific cyclization chemistries to be removed from consideration, which is useful when a particular cyclization is difficult to form or is prohibited. Yet it also yields insights into the effect of *P*(*C*_*i*_|*x*) (our exponential reweighting) on our loss minimization. Intuitively, the fewer allowed cyclizations, the greater the post-conditioning losses should be; fewer considered cyclizations means fewer options for the framework when annealing a given prior sample. As such, we condition DNF generation of the peptide chains used in Fig. 3.1, albeit restricted to just head-to-tail amide bonds, as this cyclization is common to all peptides. This provides a useful sanity check: if this distribution of H2T-restricted ℒ_cyc_ is generally greater than that of all cyclizations, this suggests our CycLOPS annealing protocol may correspond to valid cyclizations and that post annealing distributions of ℒ_cyc_ may reveal how amenable a peptide is to the considered cyclizations. If, however, the underlying Boltzmann generator is not trained well, such that it produces structures with little correspondence to valid conformations, we would expect these distributions to be somewhat similar, as optima will be unrestrained by what is physically possible. Fortunately, the expected distinction between restricted and unrestricted cyclization is observed in Fig. 10. For all tested peptides, restricting loss minimization to just H2T amide bonding significantly increases the mean of the distribution and significantly alters the distribution itself under a two-sample Kolmogorov-Smirnov test.

**Figure 10.**
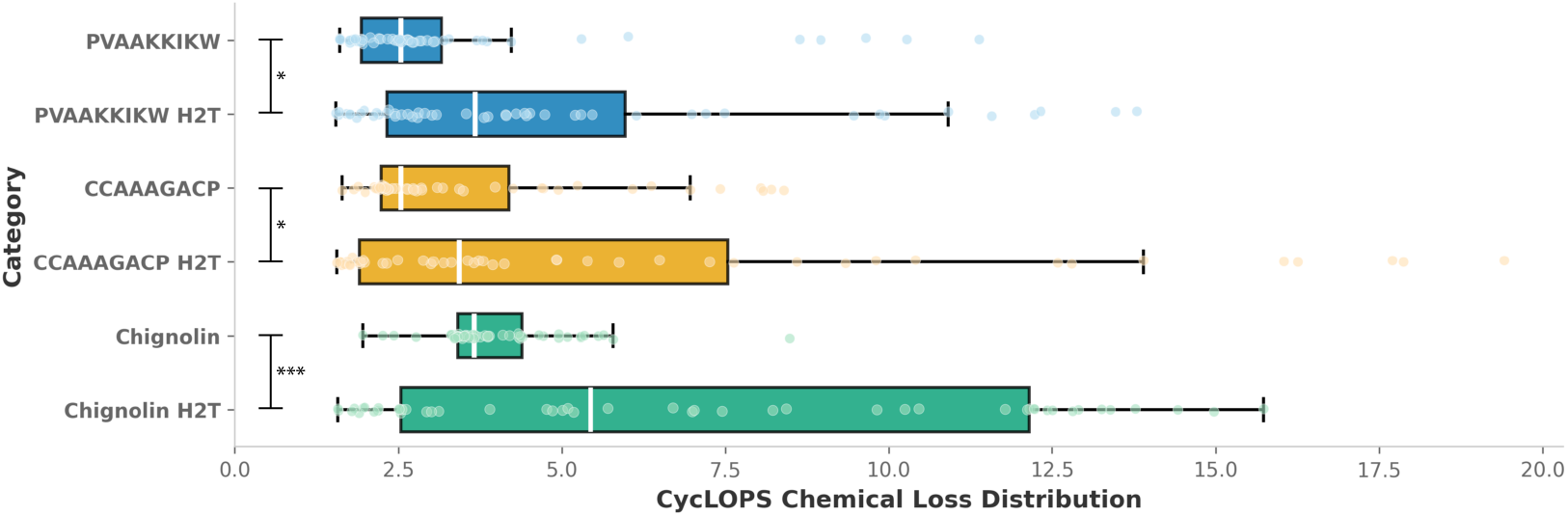
Box and whisker plot showing the distribution of CycLOPS cyclic loss values for 50 random samples of representative peptides after simulated annealing, comparing all-cyclizations (samples from Fig. 7) versus head-to-tail amide bonds only (H2T). H2T distributions show higher mean and median values and are statistically distinct from unrestricted cyclization (* *p* < 0.05, *** *p* < 0.001, two-sample Kolmogorov–Smirnov test). Interestingly, H2T distributions also exhibit greater variance for all peptides tested. All differences in means between unrestricted and H2T annealing are also significant under a permutation test with 10,000 random permutations (*p* < 0.05).

While ablation and simulated annealing hyperparameter studies are left to future work, we do include a brief examination of the latent dynamics during the optimization process in Sec. B.1. This reveals an exciting direction for the improvement of CycLOPS

### 3.2 Conditional Normalizing Flow (ECNF++) Flow Guidance

We generally observe that, for the peptides tested, ECNF++ guided flow produces less satisfactory conformational samples than SBG simulated annealing. Of particular significance is that the peptides employed in this study are limited to those permitting only head-to-tail amide bond cyclization. This constraint arises from the scarcity of suitable MD trajectories in the existing literature—a limitation that is further exacerbated by the exponential scaling of inference time with system size for state-of-the-art CNF-based Boltzmann generators. Consequently, only small peptides are currently computationally feasible, which severely restricts the already limited pool of available training data. Moreover, as loss guidance is inherently based on gradient descent, we suspect it may be more prone to getting trapped in local minima than simulated annealing, as it lacks the explicit exploratory potential of the latter.

However, our flow guidance succeeds in significantly reducing distribution of CycLOPS for all tested peptides, as shown in Fig. 11, though not to the degree of the simulated annealing shown in Fig. 7. In all cases, this increases the variance of the distributions, and produces apparent clusters of datapoints at the lower extrema of our losses. It should be noted that since the ECNF++ architecture is E(3) equivariant, it is prone to producing samples of incorrect chirality or mixed correct-incorrect chirality. Samples of entirely wrong chirality are mirrored and included, whilst those of mixed chirality are discarded. For the subsequent analyses, we begin with 128 unconditional and conditional samples and filter based on the aforementioned procedure.

**Figure 11.**
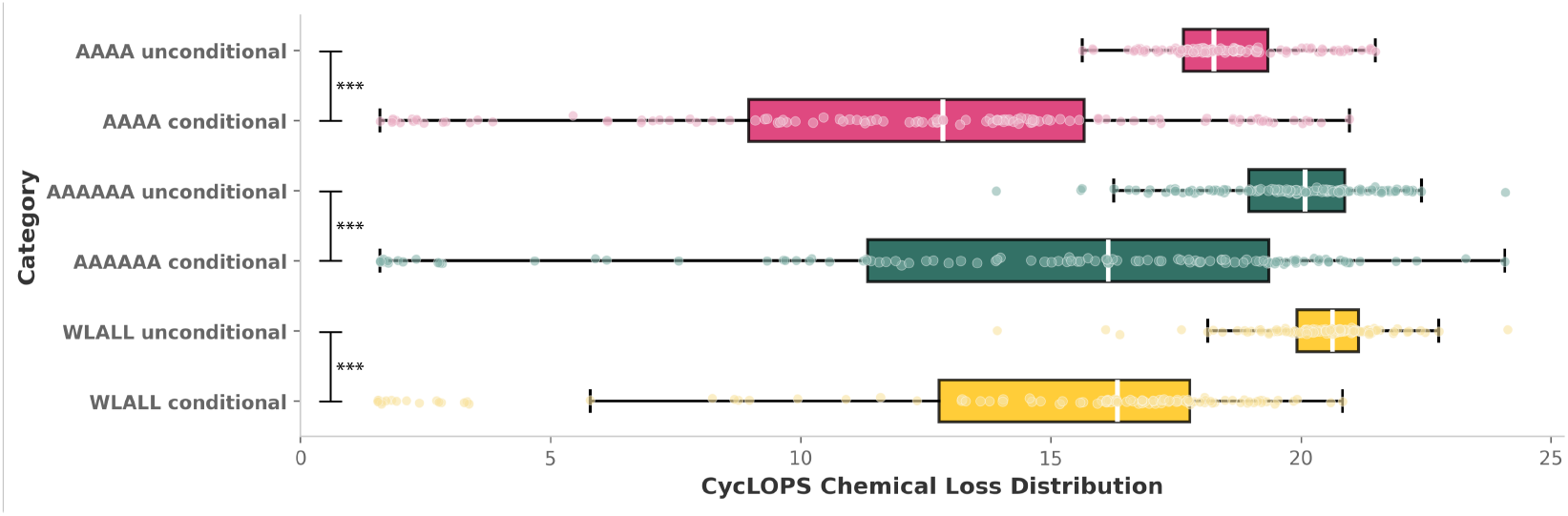
Box and whisker plots showing the distribution of CycLOPS loss values for unconditioned and conditioned-with-guided-flow ECNF++ samples across various peptide sequences. As the ECNF++ has E(3) equivariance, samples of entirely the wrong chirality are reflected, whilst those of mixed chirality are discarded. All conditioning was performed using loss-flow guidance with ω = 1.06. For each sequence, conditioning significantly alters the loss distribution (two-sample Kolmogorov–Smirnov test) and significantly reduces the mean loss (permutation test with 10,000 random permutations), with *p* < 0.005 for both tests.

We examine the apparent minimum clusters on a sample-wise level by visualizing the conformation of the four conditioned samples with the lowest CycLOPS ℒ_cyc_ loss via NGLView Nguyen et al. (2018). We then compare these to the conformation of the four unconditional samples with the lowest ℒ_cyc_ to verify that this loss reduction genuinely corresponds to increased cyclicality. We do this for three peptides, AAAA, AAAAAA and WLALL, as shown in Fig. 12. We see, therefore, that conditioning does correspond to an increase in sample cyclicality. However, this repeatedly yields low sample quality. For instance, the C-terminal cap which is added to tetra- and hexa-alanine, as a requirement of Amber MD force field, frequently “separates” from the rest of the molecule. The same occurs with the bulky sidechains of Tryptophan when conditioning, as shown in Fig. 12 C). We may therefore conclude head-to-tail amide cyclization drives the flow out of distribution. Moreover, the underlying ECNF++ network visibly struggles with the C-terminal cap even without conditioning, though this may be a consequence of the improper initialization of atom encodings – e.g. accidentally creating indistinguishable hydrogens. As such, CycLOPS-based guided flow conditioning should be further studied on peptides which admit more cyclization chemistries, particularly those which are more naturally amenable given their Boltzmann distribution - none of the present structures are intuitively very inclined toward head-to-tail amide cyclization without perturbing their equilibrium distributions due to the presence of relatively large chemical groups near the termini (see Sec:A.3). Nevertheless, CycLOPS effectively drives conformational sampling toward cyclic structures.

**Figure 12.**
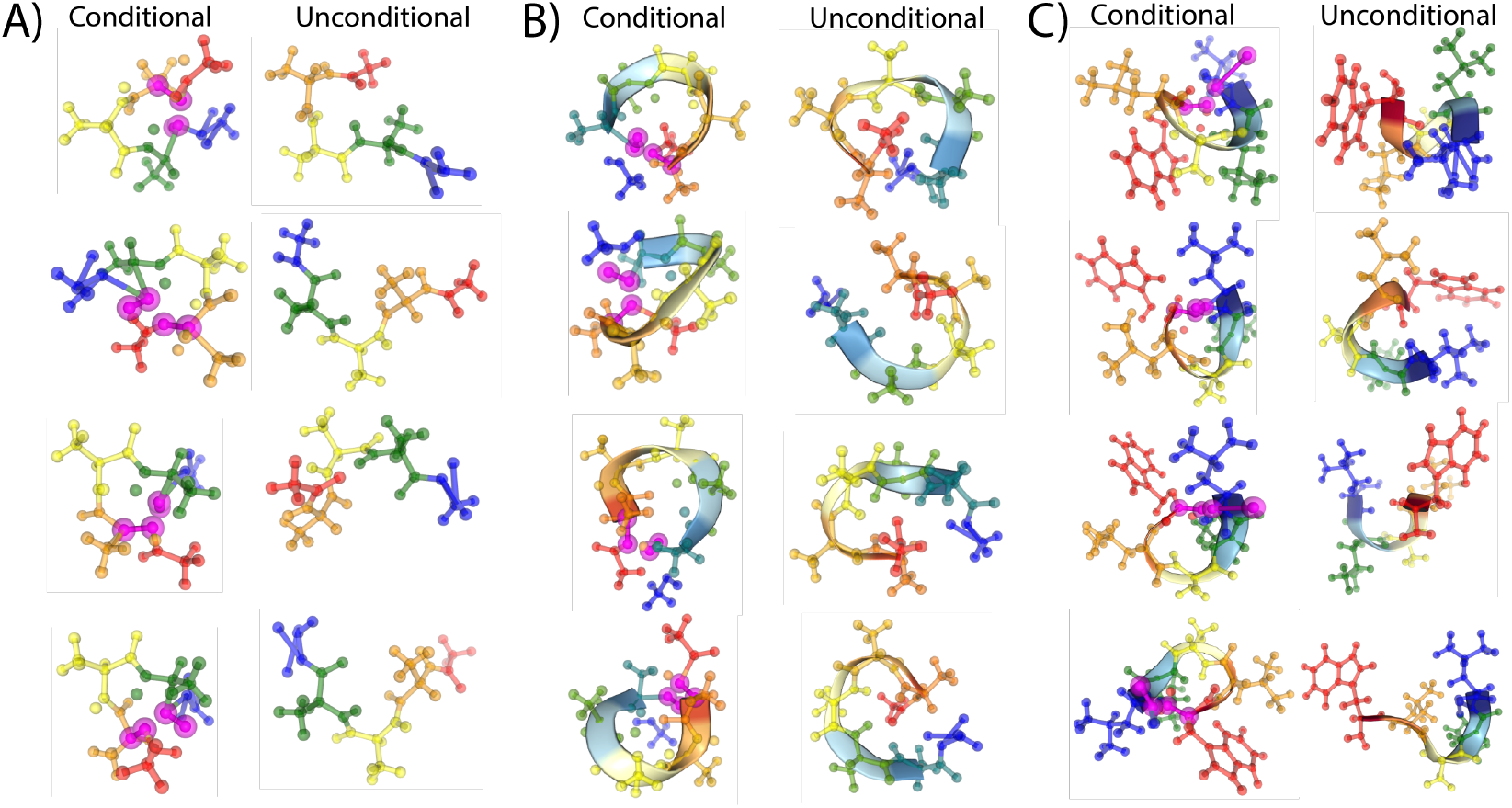
Comparison of chirality-corrected conformations generated by guided flow conditioning v.s. unconditional generation. Shown are the four lowest CycLOPS loss conformations from each method for peptide sequences: **(A)** AAAA, **(B)** AAAAAA, and **(C)** WLALL. Pink highlights indicate CycLOPS tetrahedron atoms for conditional sampling.

## 4 Conclusion and Future work

We present CycLOPS, the first model-agnostic framework that leverages linear peptide data to design cyclic structures across diverse linkages formed through complex cyclization chemical reactions. Our tetrahedral geometry approach, using KDE-fitted loss functions from toy linkage MD simulations, successfully conditions Boltzmann generators to produce valid cyclic conformations. Across multiple peptide sequences, both DNF-simulated annealing and CNF-guided conditioning achieve statistically significant mean loss reductions, with DNF approaches demonstrating superior sample quality. The framework advances beyond traditional head-to-tail and disulfide-only constraints by automatically identifying optimal cyclization strategies – from the set of 18 implemented cyclizations – through maximum entropy reweighting; this can be trivially extended to many more.

Key limitations include assumptions about small linkage chemistries and optimization parameter sensitivity (see Sec B.1). Future work should focus on: (1) expansion of the cyclization chemistries through the implementation of new MD-based cyclic losses, (2) integration of binding motif preservation with cyclization optimization, and (3) application to therapeutically relevant peptides tar-geting cancer-associated proteins (PD-1, MDM2, MDMX) and enzymes, and (4) conditioning sequence-transferable Boltzmann generators when available. Validation through molecular docking and experimental binding assays will be critical for establishing CycLOPS utility in drug discovery.

## Acknowledgments

The authors would like to thank Charlie Tan, Amalie Gudum, Klavs Jermakovs, Matthew Greenig, Sebastian Pujalte Ojeda, Karla Milcic, and Chaitanya K. Joshi for useful comments and insights. The training and sampling of Boltzmann generators was made on the Cambridge CSD3 cluster.

## >A Appendix: Additional Theory and Setup

### A.1 Proof of Maximum Entropy

By Bayes’ rule

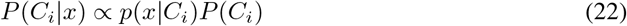

If one has a plethora of MD trajectories for diverse chains, one could compute 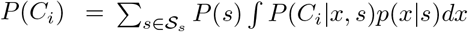 where *S*_*s*_ is the set of all amino acid sequences. As there is no such data widely available yet, we make the maximum entropy assumption that *P*(*C*_*i*_) is uniform over its support, *P*(*C*_*i*_) ∝ 1, yielding:

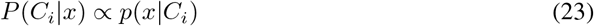

### A.2 Batch Simulated Annealing Algorithm

We present the algorithm used to perform latent space optimization. In practice, we set temperature determining κ to 2, cooling rate λ to 0.995, and define our objective function *E*(·) ≡ ℒ_cyc_(·). It may be advantageous to construct the objective function more cleverly, such that it considers additional desiderata. This may include binding motif preservation or OpenMM-based conformational energy.

#### Algorithm 2

Batch Simulated Annealing for Chemical Loss Optimization

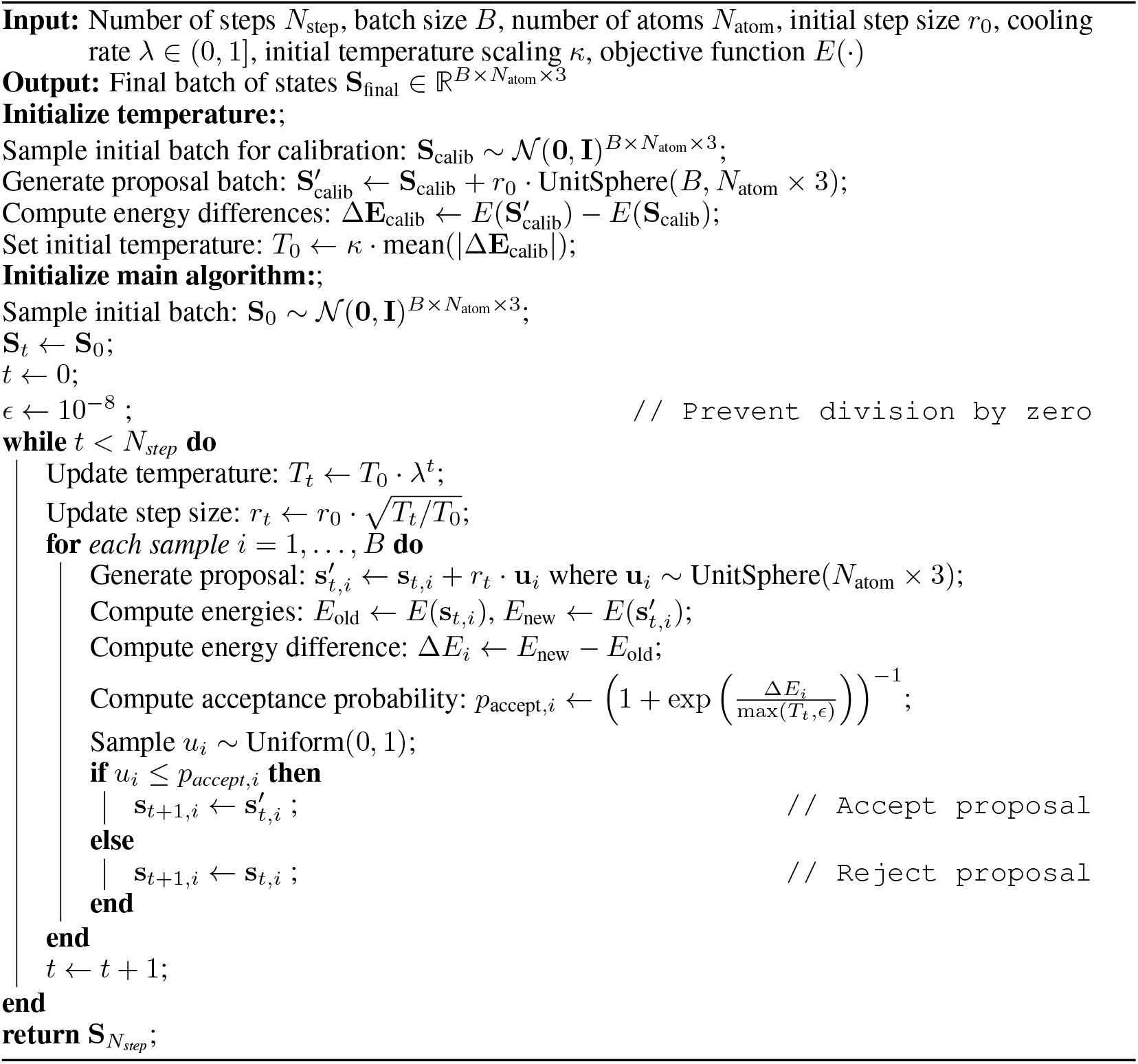

### A.3 Guided Flow Details

Following the empirical approach suggested by Komorowska et al. (2025), we set the Jacobian 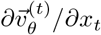 to **0**, bypassing this computationally expensive calculation. We additionally find that generated samples are quite sensitive to our choice of ω, modulating quickly between linear and cyclic conformational space, but at the cost of sample quality. All included sequences only permit head-to-tail cyclizations out of those considered by CycLOPS currently, and tetra- and hexa-alanine contain N- and C-terminal caps from AmberTools MD simulations. Furthermore, the bulky tryptophan sidechain appears to hinder head-to-tail linkage conditioning, potentially driving predicted structures well outside the training distribution. Regardless, we use ω = 1.06 for all ECNF++ conditioning.

## B Appendix: Additional Experimental Results

### B.1 Oss minimization during simulated annealing

Examining the optimization dynamics reveals an important consideration. Fig. 13 displays the progression of ℒ_cyc_ values throughout the 2000-step simulated annealing process for both unrestricted and head-to-tail restricted cyclizations. Notably, the loss values appear to plateau in the latter stages of optimization. This either suggests that the simulation may be (1) converging rather quickly, and hence cooling too fast, or (2) that more steps than necessary are used during the optimization. Given Fig. 8 and Fig. 9, however, we suspect latent space is at least somewhat well explored. Still, this suggests the need of thorough studies on the effects of the various simulated annealing parameters on convergence and cyclization choice, which we leave for future work.

**Figure 13.**
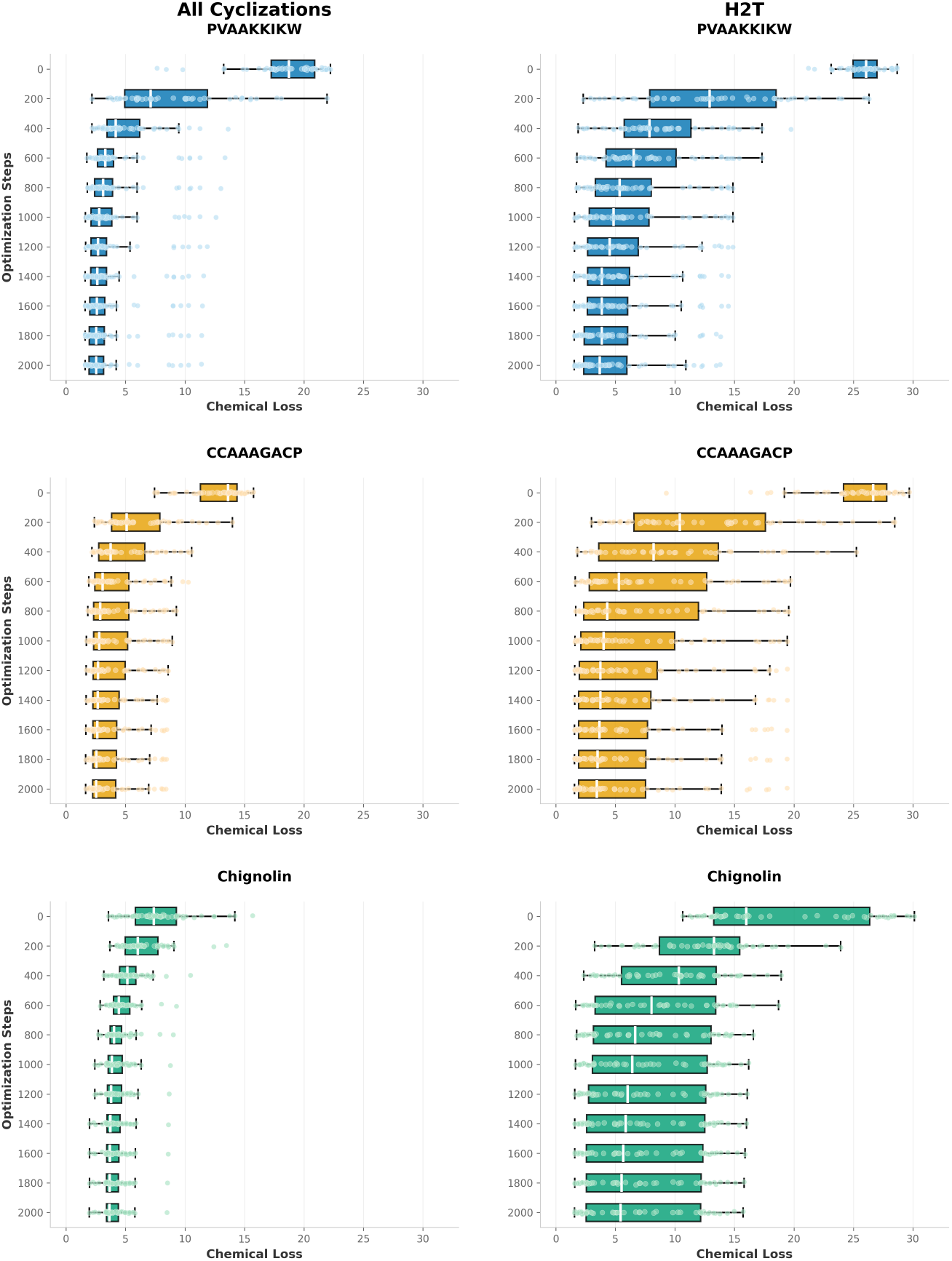
Box and whisker plots of CycLOPS cyclic loss over 2000 steps of simulated annealing for peptide sequences from Fig. 3.1(rows) for all cyclizations (left column) and just head-to-tail (right column, H2T).

### B.2 Computational Resources of Model Training

Training the Boltzmann generators that underlie CycLOPS requires significant computational resources. We include these here, to highlight the importance of conditioning methodologies which do not require retraining.

## Footnotes

1 Jiang et al. (2025) ^consider a similar formulation, though *p*(*x*) is implicitly learned during training.^

2 Sufficiently small can be defined as small enough that atoms outside of this tetrahedron do not interact with the linkage’s virtual atoms. This represents a significant limitation.

3 Of course, this assumes 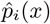is unaffected by the change. This is valid, because *T* represents a *virtual* as opposed to physical temperature; it is essentially just a coefficient.

4 This is only rigorously true if the diffusion uses DDIM sampling (Song et al., 2020) and the flow matching ODE is solved with Eulerian integration. We direct the reader to Gao et al. (2024) for more details.

